# Position Effect Variegation in natural populations not explained by common variation in known modifiers

**DOI:** 10.1101/129999

**Authors:** Keegan J. P. Kelsey, Andrew G. Clark

## Abstract

Changes in chromatin state may drive changes in gene expression, and it is of growing interest to understand the population genetic forces that drive differences in chromatin state. Here, we use the phenomenon of position effect variegation (PEV), a well-studied proxy for chromatin state, to explore the genetic architecture of natural variation in factors that modify chromatin state. While previous mutation screens have identified over 150 suppressors and enhancers of PEV, it remains unknown to what extent allelic variation in these modifiers mediates inter-individual variation in chromatin state. Is natural variation in PEV mediated by segregating variation in known Su(var) and E(var) genes, or is the trait polygenic, with many variants mapping elsewhere in the genome? We designed a mapping study that directly answers this question and suggests that the bulk of the variance in PEV does not map to genes with prior annotated impact to PEV. Instead, we find enrichment of top *P*-value ranked associations that suggest impact to active promoter and transcription start site proximal regions. This work provides a quantitative view of the role naturally segregating autosomal variants play in modifying PEV, a phenomenon that continues to shape our understanding of chromatin state and epigenetics.

---

**C**hromatin states, through their impact on chromatin accessibility and gene expression have clear evolutionary importance (Schulze and Wallrath 2007). The ModEncode project has generated valuable data for use in understanding determinants of chromatin states (modENCODE Consortium *et al.* 2010). This has generated genome-wide assessment of chromatin features and led to identification of broad chromatin features with functional consequences (Filion *et al.* 2016; Kharchenko *et al.* 2011; Ernst and Kellis 2012). Despite rapid progress in chromatin biology, we still know little about naturally segregating variants and involvement in generating differences in chromatin states among individuals. Any natural variant that serves to impact chromatin state may be a target of natural selection, and it is of great interest to understand population genetic forces that drive differences between chromatin features of individuals.

Originally discovered by H. J. Muller, PEV has long been studied in *Drosophila melanogaster* and is widely accepted as a valuable tool in understanding dynamics of chromatin state and gene expression, especially with regards to the boundary between heterochromatin and euchromatin. The most commonly used form of PEV is the result of a specific X chromosome inversion, *white*^*mottled-4*^ or simply *w*^*m4*^ (Muller 1930), that relocated the normally euchromatic *white* gene next to pericentromeric heterochromatin. The molecular impact of this relocation is now understood to be a spreading of chromatin factors that results in altered chromatin structure and gene silencing of *white* without known modification to coding sequence (Wallrath and Elgin 1995). Depending on the genomic background, the phenotypic manifestation of this particular silencing is mosaic across the facets of the eye, resulting in a “mottled” eye phenotype with local clones of cells in each eye showing an apparently stochastic response. Of significance, PEV is triggered by gene-chromosome rearrangements in many organisms, producing modified gene expression as a result of modified chromatin environment (Girton and Johansen 2008; Elgin and Reuter 2013).

Manifestation of PEV in the *Drosophila* eye allows for ease of visible phenotyping and has generated a large body of work leading to the discovery of numerous PEV modifying factors, both genetic and environmental, that also impact chromatin states. These modifiers are typically described as positively influencing gene expression at the *w*^*m4*^ locus, termed suppressor of variegation, or Su(var), or negatively influencing gene expression at *w*^*m4*^, termed enhancer of variegation, or E(var). Screens for modifiers have identified over 150 loci that modify PEV (Ebert *et al.* 2004). Two such well characterized modifiers of PEV and chromatin include *Su(var)3-9* and *Su(var)2-5*. *Su(var)3-9*, a histone methyltransferase with action on histone H3 at lysine 9, helps to form heterochromatin (Schotta *et al.* 2002). *Su(var)2-5* or *HP1a* interacts with *Su(var)3-9* to uniquely bind distinct regions of the genome (Greil *et al.* 2003) and is required for the spread of heterochromatin (Hines *et al.* 2009). It is important to understand, however, that the vast majority of these modifiers have been identified through mutation screens and may or may not represent loci harboring natural variation that impact the PEV phenotype. Despite the extensive literature on modifiers of PEV, there is currently little information on natural variation and impact to chromatin state between individuals and populations.

Genome-wide association studies (GWAS) initially gained favor in human studies as a way to perform a relatively unbiased search for common natural variants involved in a phenotype of interest. Using sets of inbred reference lines of *D. melanogaster*, GWAS has proven to be a powerful resource for generating unbiased, data-driven hypotheses, where a genome-wide search of candidate loci may be coupled with the extensive functional annotation and genetic tools already available. The *Drosophila* Genetic Reference Panel (DGRP), a collection of genome-sequenced inbred lines of *D. melanogaster* (Mackay *et al.* 2012; Huang *et al.* 2014), has become a handy resource for initial GWAS screens with Drosophila. Several groups have already successfully used the resource to identify novel variants involved in a wide range of traits, including; sleep (Harbison *et al.* 2013), leg development (Grubbs *et al.* 2013), sperm competition (Chow *et al.* 2012), host-microbiota interaction (Dobson *et al.* 2015), fecundity and fitness (Durham *et al.* 2014) and nutritional indices (Unckless *et al.* 2015).

To better understand the underlying genetic architecture of segregating natural variants involved in heterochromatin dynamics, we performed GWAS on F1 progeny of DGRP lines crossed to *w*^*m4*^, a line bearing an X-linked inversion that displays PEV of the white eye phenotype. PEV was quantified by novel digital image analysis of visible images captured with a dissecting microscope. Despite this detailed work, we find little evidence of association for segregating variants within known Su(var) and E(var) genes. However, we do find variants having association with PEV to be over-represented in regions having a chromatin state indicative of active promoter and transcription start site (TSS)-proximal features. Furthermore, a comprehensive search across binding sites for factors that modify chromatin state link numerous binding sites to PEV-associated variants, emphasizing regions with bi-stable chromatin states. Altogether, the evidence suggests autosomal natural variation interacts with PEV through numerous, small effect loci that are enriched for TF binding and sites of open chromatin, implying influence through gene expression or subtle changes to chromatin balance.

## Materials and Methods

### Drosophila stocks

Lines from the *Drosophila* Genetic Reference Panel (DGRP) (Mackay *et al.* 2012) were a gift of the Mackay lab. 1712 (Bloomington), which harbors the *white*^*mottled-4*^, *In(1)w*^*m4*^ or simply, *w*^*m4*^, locus on the X chromosome and a second chromosome deletion and balancer, *Df(2L)2802*/*CyO*, was used to assess variegation across the DGRP population. Canton-S and mutant eye color stocks, 245 (*bw*^*1*^) and 3605 (*w*^*1118*^) (Bloomington) provided biological context in our eye color phenotype assay. All flies were maintained on a standard cornmeal-molasses-sucrose-yeast media and kept at 25°C on a 12-hour light/dark cycle.

### Experimental cross

In each of two replicate vials, ten males from each DGRP line were crossed to three virgin females of the 1712 stock and allowed to mate for one day. Mated 1712 females were then transferred to new food, allowed to lay eggs, and removed from the vial after five days. Variegating male progeny segregated into two phenotypic classes based on the second chromosome, curly winged (*CyO*) and non-curly winged (*Df(2L)2802*), and were aged at least four to eight days before imaging.

### Image capture and eye color quantification

The left or right eye was randomly selected from each adult male and imaged using an Olympus SMZ-10 dissecting microscope with an attached Cannon Rebel 6 megapixel digital camera in a windowless room. Images were captured and stored using software from the camera manufacturer. On average, 17.5 eyes (standard deviation = 9.2) were imaged from each line, resulting in a total of 3966 images across all lines and second chromosome combinations. A standard gray card (Kodak, 18% gray) was imaged before and after each set of conditions to normalize against fluctuating light conditions. Ommatidia from images were isolated using a pipeline built in Cell Profiler 2.0 (Lamprecht *et al.* 2007) and visually inspected. Images that failed to process were individually assessed and isolated in Photoshop. Isolated ommatidia and standard gray card images were then processed using custom scripts in R (R Development Core Team 2008). Ommatidia image files were separated into red, green, and blue color channels and values were bounded between 0 and 1. Color channel values from each pixel of every image were normalized against the mean value of individual color channels from matched gray card pairs according to a generalized gamma adjustment using the formula,

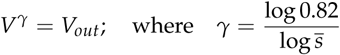

where 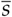 is the individual mean color channel (red, green or blue) for the gray card imaged before and after samples. 0.82 refers to the idealized 18% gray card value in RGB color space. *V* is the color channel value for an individual pixel within the image, *γ* is the normalizing function, and *V*_out_ is the normalized color value. The final summarized output for each eye and image resulted in three values, including the mean of each normalized red, green and blue color channel. We show that these descriptive values sufficiently separate individuals for our purposes.

### Statistical analysis of phenotype

All subsequent analysis was performed in R (R Development Core Team 2008). The mean red, green and blue color values for each image were used input for Principal Component Analysis (PCA). Eye groups were assessed using MANOVA and the formula,

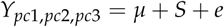

where *Y* is PC1, PC2 and PC3 from each image, and *S* is the stock of origin (Canton-S, 1712, 245 or 3605). Differences between experimental groups were assessed using a linear model to fit the principal components from each image with the formula,

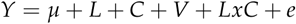

where *Y* is either PC1, PC2 or PC3, *L* is the DGRP line of origin, *C* is the second chromosome background (*CyO* or *Df(2L)2802*), and *V* is the replicate vial (A or B). Proportion of variance for each effect is calculated as,

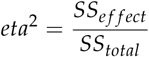

where *SS* is the sum of squares. Broad-sense heritability (H^2^) was calculated using,

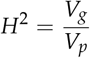

and based on the linear model,

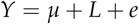

where *Y* is a single principal component and single second chromosome background combination, and *L* is the DGRP line of origin. The proportion of variance explained by line differences was considered the variance attributed to genetic components (*V*_*g*_), and the total variance of the sample was considered the variance attributed to phenotype (*V*_*p*_). For each second chromosome background, H^2^ was summed across the three principal components and weighted according to the proportion of variance explained by each PC (Table S1).

### Genotypes and association testing

Genotypes and annotation for DGRP lines were downloaded from the website, dgrp.gnets.ncsu.edu, and all variants and findings are reported using build BDGPR5/dm3. As others have observed, the DGRP lines display small amounts of cryptic genetic relatedness (Huang *et al.* 2014; He *et al.* 2014). Here, we used GEMMA (Zhou and Stephens 2012) to both estimate a centered genetic relatedness matrix (GRM), accounting for cryptic relatedness, and implement the univariate mixed linear model (MLM). PC1 from each of the two experimental populations (*CyO* and *Df(2L)2802* second chromosome backgrounds) was used as separate input, providing two independent sources of association values. Males were regressed on each variant using single marker association (SMA). Variants were treated as a fixed effect and the GRM was included as a random effect. Effect size was determined using the function, pes, within the R package compute.es (Del Re 2013). 107 lines were used with a *CyO* second chromosome background and 109 lines were used with a *Df(2L)2802* second chromosome background. Testing was performed across 775,689 (*CyO*) and 928,587 (*Df(2L)2802*) bi-allelic variants (SNPs and indels) with a MAF of 0.05 or greater. Due to the high correlation between phenotype and top associations of the two experimental populations, subsequent analysis was performed using only the *Df(2L)2802* GWA data. Bootstrap analysis was used to generated expected site class frequencies. Site classes were counted from 1,000 randomly selected common variants, and an expected distribution was achieved through 10,000 iterations of resampling. For variants with more than one annotated class, only one class was selected and priority was ranked as follows; nonsynonymous > ncRNA > synonymous > UTR > intronic > intergenic. The prop.test in R was used to assess the proportion of exonic variants that were nonsynonymous between observed counts in the variants with top associations and expected counts through sampling.

### Candidate gene analysis

Candidate genes were selected based on known involvement in chromatin modification. Genes were identified in Flybase using the term “Modifier of Variegation.” Location of each gene was extracted from Flybase and SNPs within the gene and +/-2 kb of the gene were examined using the above mixed linear model for each SNP. Bootstrap analysis was used to generate an expected Site Frequency Spectrum. Proportions of variant frequencies were tallied by selecting 1,000 variants at random and then binning into allele frequency groups of 0.05. An expected distribution was achieved through 10,000 iterations of resampling. Bootstrap analysis was used to generate an expected distribution of segregating sites with sets of genes. Total number of segregating sites (within ±2 kb) were counted within sets of 105 randomly selected genes throughout the autosomal genome and normalized by number of base pairs summed across the full set. The expected distribution of proportion of segregating variants was then achieved through 10,000 iterations of resampling.

### Genomic analysis

The 9-state genome-wide combinatorial chromatin state annotation is described in Kharchenko *et al.* (2011) and annotation files were sourced from www.modencode.org. Expected 9-state distributions were generated in the experimental population through sampling autosomal variants for state assignment. 1,000 variants were randomly selected and chromatin states were counted. This process was repeated 100 times. ChIP-chip and ChIP-seq files were also downloaded from ModENCODE (http://www.modencode.org). If replicate samples existed, only one file was randomly selected for analysis and composite files, if they were made available, were used instead of individual samples. Comparison between the expected distribution of variants within binding sites and variants enriched with associations to PEV was performed as described above. 1,000 autosomal variants were selected at random and variants within binding sites were counted. The distribution of counts, as generated through 10,000 iterations, was then compared to observed counts from 1,000 of the top *P*-value ranked associating variants. A full list of factors with respective ModENCODE IDs and observed and expected counts has been made available (File S3).

### Data Availability

Original eye images are available upon request. File S1 contains PEV phenotype values. File S2 contains a full list of variants with respective association *P*-values. File S3 contains observed and expected counts of top PEV associations within annotated chromatin features.

## Results

### Image analysis captures multidimensional color space

Pigments in the eye of *D. melanogaster* are synthesized by two, well-characterized metabolic pathways (Summers *et al.* 1982). These pigments are typically quantified through separate extractions based on chemical properties (Ephrussi and Herold 1944). Often, only a single extracted pigment is used to describe eye color, and, although useful for detecting general differences, this method results in considerable loss of information. Even casual inspection of eye color patterns that manifest PEV reveals a far more complex range of differences, including pigment intensity, different hues of pigmentation including yellow, orange, brown and red, and variation in patch size and morphology. To better capture the multidimensional aspects of eye color and PEV, and improve mapping, we developed an imaging method that retains and fully describes eye color within a single assay (see methods). Eye-color stocks (Figure 1A) were imaged and Principal Component Analysis (PCA) using the values from the individual red, green and blue color channels, of each image, sufficiently described the multivariate data. Non-pigmented (*white*) and pigmented stocks (*bw*^*1*^, *w*^*m4*^, and Canton-S) exhibited large differences described by principal component 1 (PC1), and pigmented stocks (*bw*^*1*^, *w*^*m4*^, and Canton-S) exhibited differences primarily described across principal component 2 (PC2) (Figure 1B). PC1 accounted for 87.4% (s.d. = 24.4%) of variance in the data and PC2 accounted for 12.5% (s.d. = 9.2%). Component loadings provide detail on how each color channel contributes to the dispersion of the data, where blue and green channels have a similar impact on PC1 (−0.72 and −0.68, respectively) while the red channel has a minor impact (−0.12). This is in contrast to PC2, where the red channel is the primary driver (0.94) and blue and green channels have minor roles (−0.30 and 0.15, respectively). MANOVA using PC1 and PC2 from each individual image highlights the ability of this approach to discriminate across eye groups of the four stocks (*P*-value < 2.2×10^−16^).

**Figure 1.**
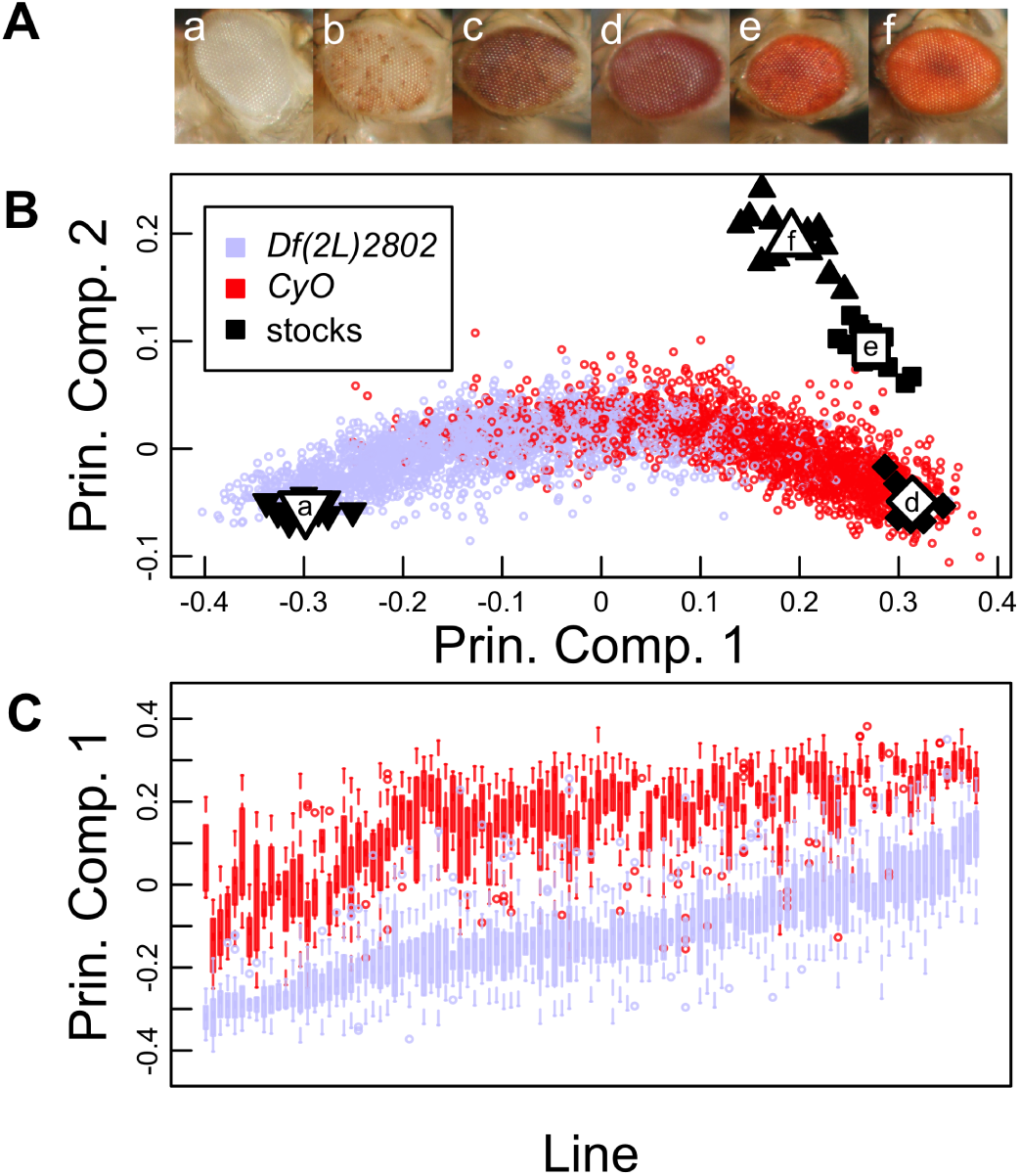
**(A)** Examples of imaged eyes from various *Drosophila melanogaster* alleles; (a) *w*^*1118*^, (b) *w*^*m4*^ with the *Df(2L)2802* second chromosome, (c) *w*^*m4*^ with the *CyO* second chromosome, (d) *bw*^*1*^, (e) *w*^*m4*^ with *Df(2L)2802*/*CyO*, and (f) Canton-S. **(B)** Scatter plot of PC1 and PC2 values for mutant and experimental PEV eyes. Black and white points represent individual images and average values of eye colors from stocks; (a) *w*^*m4*^, (d) *bw*^*1*^, (e) *w*^*m4*^ with *Df(2L)2802*/*CyO*, (f) Canton-S. Experimental individuals with just the *Df(2L)2802* (periwinkle), in general, show greater variegation and eyes that are closer to the *w*^*1118*^ allele (a), an eye that lacks pigmentation. Experimental siblings with the *CyO* allele (red) show less variegation, or more pigmentation. **(C)** Boxplot of PEV summarized by line (*x*-axis) and separated by second chromosome, *Df(2L)2802* (periwinkle) vs. *CyO* (red). PEV is represented PC1 (*y*-axis), where lower values indicate less pigmented eyes and higher values indicate more pigmented eyes. Among-line variance is greater than within-line variance, suggesting natural genetic variation is involved in observed differences in PEV.

### Natural variation in background genetic effects on white-mottled-4 expression

To quantify natural variation in PEV, we made use of the *Drosophila* Genetic Reference Panel (DGRP). The PEV phenotype was expressed by crossing males from inbred DGRP lines to virgin females carrying the *white-mottled-4* (*w*^*m4*^) allele on the X chromosome. F1 variegating males were identical across a single X, third and fourth chromosome, segregating according to one of two second chromosomes and varying with respect to a full haplotype from each of the DGRP lines assayed (Figure S1). The two second chromosomes differ primarily with respect to an approximate 200 kb deletion in 25F2-25F5 (*Df(2L)2802*) on the non-balancing chromosome and an inversion on the balancing chromosome (*CyO*). Experimental F1 progeny exhibited a wide range of eye pigmentation differences, showing variation that spanned a complete lack of pigmentation to eyes that were heavily pigmented (Figure S2). The quantitative image assay further detailed a broad phenotypic spread in PEV with PC1 and PC2 values spanning between an eye mutant that lacks pigmentation (*white*) and mutants of known pigment deficiencies (Figure 1B). Reapplying PCA to just the mean red, green and blue color values of images from each of the F1 variegation males, indicates that PC1 captures the vast majority of variance in the experimental data, 97.4% (s.d. = 18.6%), while PC2 and PC3 only describe a small proportion of variance, 2.4% (s.d. = 2.9%) and 0.2% (s.d. = 0.9%). Using PCA on color images of PEV individuals, effectively allows the simplification of a multivariate data source to a single describing variable, PC1, with minimal, 2.6%, loss of data and provides a robust univariate phenotype for association mapping.

ANOVA using individual PCs provides an assessment of importance ascribed to each of three experimental variables; among line differences, second chromosome differences and replicate environments (Table 1). Adjusted values show that combined genetic components explain 79.6% of the phenotypic variance, where second chromosome differences separately explained 49.5% of the variance, among-line variation (our source of natural variation) explained 27.4% of the variance and 2.7% of the variance was explained through genetic interactions of second chromosome background and individual lines. Partitioning the sample by presence/absence of the second chromosome deficiency provides two separate measures of broad-sense heritability (H^2^) given different genetic backgrounds. Among-line differences explained over half of the phenotypic variance, 59.4% and 57.4%, for each of the populations with *CyO* and *Df(2L)2802* second chromosome backgrounds (Table S1). These data suggest that the bulk of the observed variation in PEV is attributed to segregating genetic variants among the naturally derived DGRP haplotypes.

**Table 1.** Partitioning the variance in PEV attributed to genetic and environmental factors.

| Variable | Principal Component 1 |  | Principal Component 2 |  | Adjusted<sup>a</sup> |
| --- | --- | --- | --- | --- | --- |
|  | Proportion of variance | *P*-value | Proportion of variance | *P*-value | Proportion of variance |
| Among line | 27.7% | $< 1.0 \times 10^{-22}$ | 15.3% | $< 1.0 \times 10^{-22}$ | 27.4% |
| Second chromosome | 50.9% | $< 1.0 \times 10^{-22}$ | $< 0.1\%$ | 0.88 | 49.5% |
| Vial | $< 0.1\%$ | 0.33 | $< 0.33\%$ | $1.6 \times 10^{-6}$ | $< 0.1\%$ |
| Among line x second chromosome | 1.9% | $< 1.0 \times 10^{-22}$ | 31.2% | $< 1.0 \times 10^{-22}$ | 2.7% |
| Within line and Residuals | 19.5% | - | 53.2% | - | 20.4% |

<sup>a</sup> Adjusted values are calculated by summing across each specific variable as weighted by the proportion of the variance explained by each PC; 97.4% (PC1), 2.4% (PC2) and 0.2% (PC3). PC3 is not shown above as it explains a negligible amount of the variance in the data.

When comparing second chromosome backgrounds, lines showed strong positive correlation in PEV between PC1 values (Pearson correlation of 0.84), where the *Df(2L)2802* second chromosome acts a clear E(var) with respect to the *CyO* second chromosome balancer; showing greater variegation, or less pigmented eyes, in nearly all lines (Figure S3). Although not the focus of this study, it is important to recognize that individuals differing only by second chromosome accounted for almost half (49.5%) of the phenotypic variance observed, considerably more than explained through natural variation. The consequences of this highlight two potential scenarios. First is the possibility of an unannotated mutation in either a Su(var) or E(var) between the second chromosomes. A second possibility is that the totality of the deletion on *Df(2L)2802*, an approximate 200 kb deletion, acts to enhance variegation through a sponge or sink model, similar to what is proposed to occur with the Y chromosome (Francisco and Lemos 2014); suggesting general amounts of DNA (or chromatin) can act to suppress or enhance the PEV phenotype. We cannot distinguish these possibilities.

### Genome-wide Association testing

To assess the contribution of segregating variants to PEV, single marker analysis (SMA) was performed genome-wide using variegating F1 males from crosses between DGRP lines and the *w*^*m4*^ reporter. Full haplotypes from each of the distinct DGRP lines provided source variation for association mapping. PC1 from each of the two second chromosome populations were used as separate input into a univariate mixed linear model (MLM) accounting for cryptic relatedness (He *et al.* 2014). Testing was performed across common bi-allelic variants (SNPs and indels, MAF ≥ 0.05) using haplotypes extracted from DGRP chromosomes 2, 3 and 4. The X chromosome, carrying the *w*^*m4*^ reporter, was invariant across experimental populations. Quantile-quantile (Q-Q) plots indicate *P*-values from each experimental population overall conform well to the null distribution (Figure S4). Effect sizes follow a trend where Cohen’s *d* increases as MAF decreases (Figure S5). A comparison of the *P*-value rank ordered 1,000 top associations shows 75.3% overlap between the two experimental populations, consistent with a strong correlation in PEV between the two groups. The top-ranked associations are enriched for variants that are located in exons, UTRs and ncRNAs, with reduced representation within intronic and intergenic regions (Figure S6). A comparison of exonic sites further shows no significant difference between proportions of nonsynonymous variants in top associations compared to expected (proportion test, *P*-value = 1), suggesting a broad influence to functional elements of the genome and not strictly toward nonsynonymous variants. In addition, of the 10 variants with the smallest *P*-values in both sets of associations, only two variants were identified as resulting in a missense mutation and both variants showed weakened association between the independent GWAS samples. Among the smallest *P*-values, only two variants were identified as resulting in a missense mutation and both showed reduced association across independent GWAS samples (Figure S2). Despite many variants identified as having an effect size greater than or equal to 1 and with enrichment proximal to functional regions, our small sample sizes, substantial background effect on phenotype, as noted through second chromosome differences, and abundance of multiple variant classes in top associations reduce confidence in traditional functional follow-up. A full list of variants with association *P*-values has been made available (File S2)

### Common variants within known autosomal PEV modifiers fail to fully account for among-line differences

Despite low power to identify individual causal variants with high confidence, an extensive literature on known genic modifiers of PEV affords the opportunity to assess significance of ensembles of variants. To quantify the impact of naturally occurring polymorphism in known modifiers of PEV, we identified variants in Su(var) and E(var) genes. The term, “Modifier of Variegation” was searched within FlyBase, and over 200 genes satisfied this criterion. As our experimental setup resulted in individuals sharing a common X chromosome, the set of modifiers was reduced to 105 autosomal genes (Table S3). Variants within, or extending *±*2 kb of identified autosomal modifiers were grouped and results from the above SMA were used. 16,640 total variants (6,153 having MAF ≥ 0.05) were identified in autosomal modifiers of PEV within the 109 lines having the *Df(2L)2802* second chromosome background. Importantly, of common variants (MAF ≥ 0.05) in this reduced set of PEV genes, only 5 were also identified in the top 1,000 genome-wide SMA hits, making up less than 0.5% of top associated variants. These five variants hold overall *P*-value ranks of 379, 553, 772, 820 and 821, indicating that a majority of natural variants with likely impact to phenotypic variance of PEV do not reside in or near genes with known impact to PEV. GCTA was used to assess cumulative statistical explanation of phenotypic variance (Yang *et al.* 2011). Top variants identified through GWA, rank-ordered by *P*-value, explained a far greater proportion of among-line variance than variants within known genic modifiers (Figure 2 and Figure S7). The 1,000 top ranking overall variants statistically explained 93.7% of variance due to genetic differences and the 1,000 top ranking variants in known PEV genes explained considerably less at 56.0%. These sets are both compared to variants randomly drawn from the autosomal genome which explained, on average, 2.3% (standard deviation = 2.7%) of phenotypic variance.

**Figure 2.**
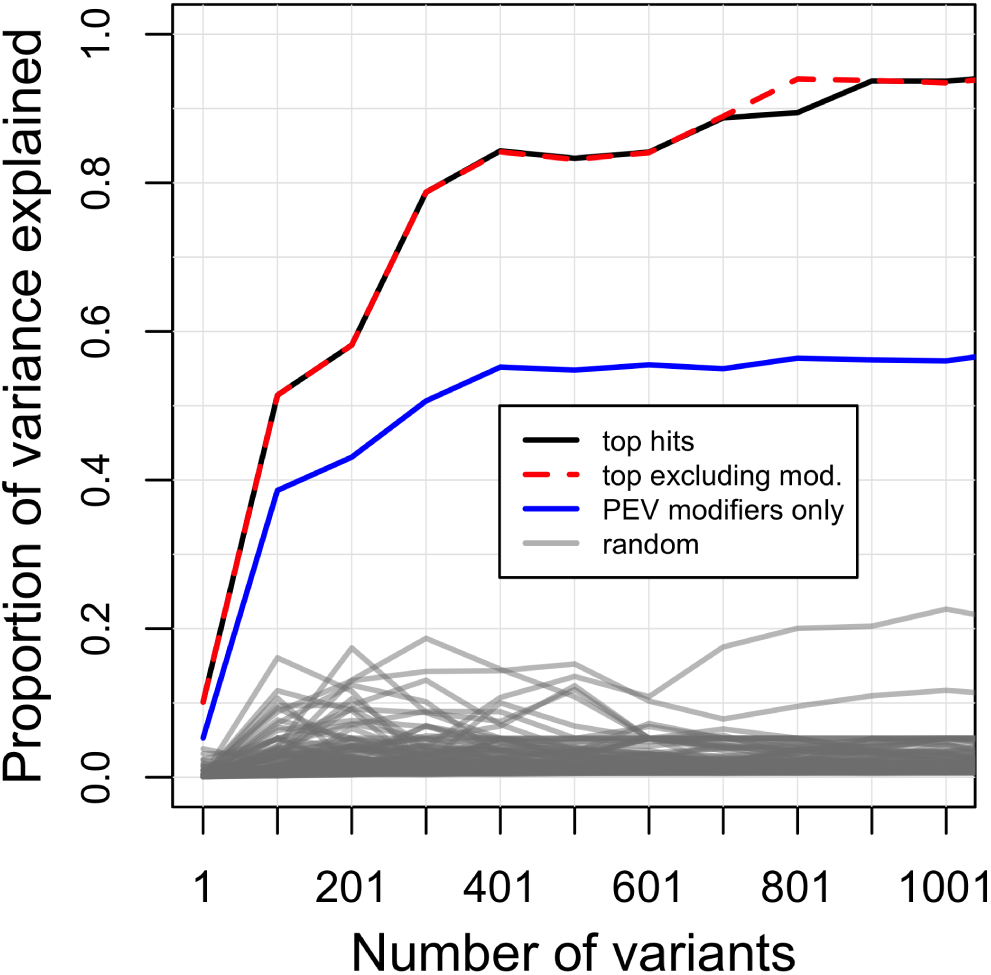
Proportion of between-line phenotypic variance explained within the *Df(2L)2802* second chromosome population using GCTA. Comparisions between SNP groupings include; the most significant GWA variants (black line), top hits excluding variants from known PEV modifiers (red line), variants within know PEV modifiers only (blue line), and randomly selected autosomal variants (gray line).

Although there is strong evidence for involvement of genes from Table S3 in PEV, there is little data indicating natural variation within these genes is responsible for differences in PEV among lines. This is not completely surprising however, as many known Su(var) and E(var) genes show conservation across species (Fodor *et al.* 2010), suggesting little room for variation in coding sequence. Indeed, two additional pieces of data further explain the lack of association with variants in known PEV modifiers and suggest purifying selection within the modifiers. The observed site frequency spectrum (SFS) within the experimental population shows an increase in low MAF variants and a decrease in variants ≥ 0.05, compared to sets of genes randomly selected from the autosomal genome (Figure 3A). Finally, known PEV modifiers exhibit a paucity of segregating sites compared to an expected distribution, having fewer segregating sites than 99.7% of gene sets randomly selected from autosomes (Figure 3B).

**Figure 3.**
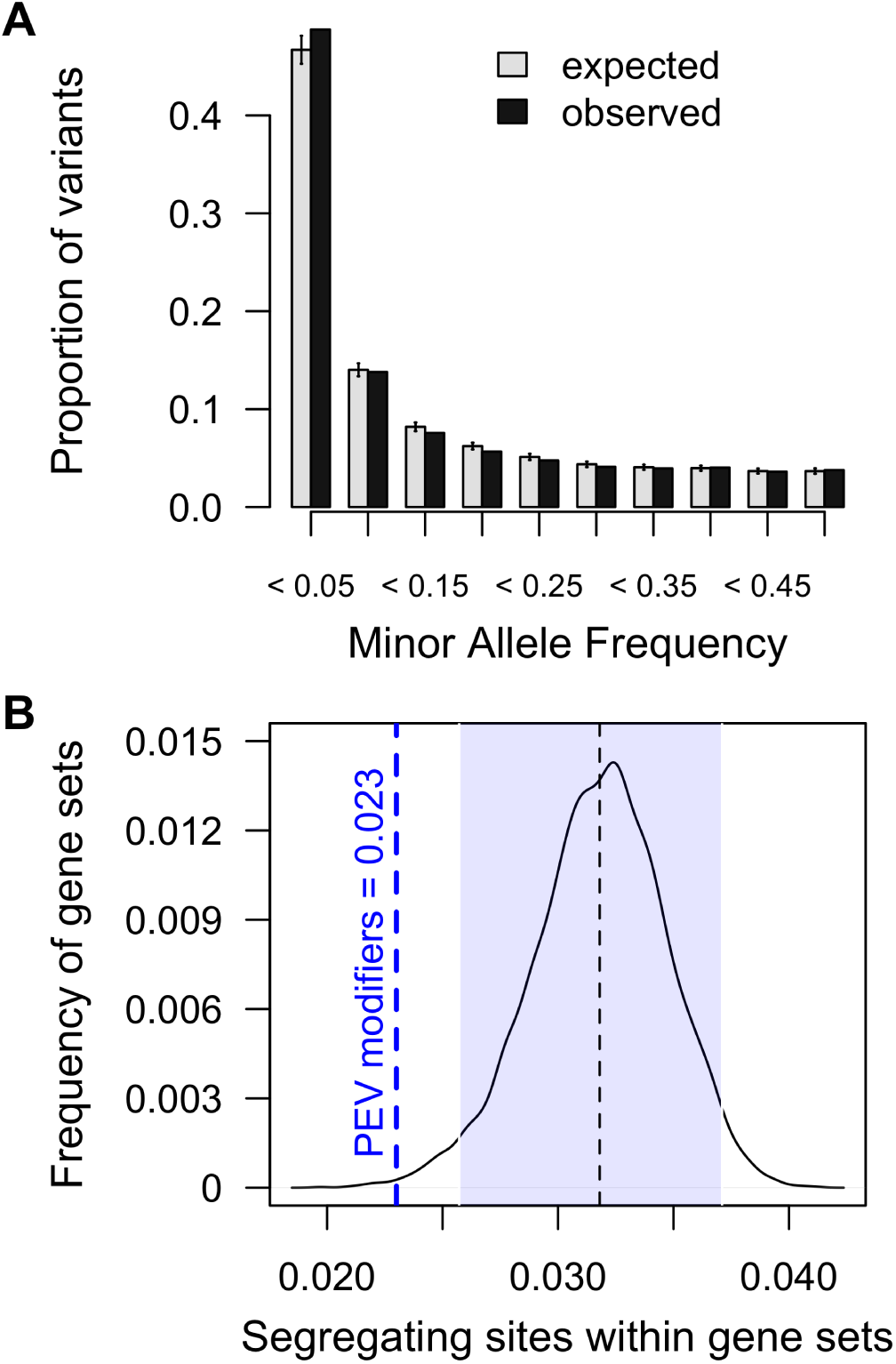
Variants within known autosomal PEV modifiers compared to expected distibutions. **(A)** Site Frequency Spectrum (SFS) of variants within known autosomal modifiers of PEV (black) and variants within sets of randomly selected autosomal genes (gray). Error bars reflect the standard deviation. **(B)** Proportion of known autosomal PEV modifiers that contain segregating sites compared to sets genes drawn randomly 10,000 times. The shaded area highlights 95% of the expected distribution. 99.7% of randomly selected gene sets contain a greater proportion of segregating sites than 105 known genic PEV modifiers.

### Chromatin and feature enrichment of top associations

As we see little evidence for association within known Su(var)s and E(var)s, we then ask if other genomic features show an over-representation of association with PEV. Chromatin features correlate with functional elements of the genome (modENCODE Consortium *et al.* 2010), and are a natural link to PEV. We next asked if particular chromatin states were over or underrepresented in the top *P*-value ranked associations. Using a previously built combinatorial 9-state (c1-c9) chromatin assignment from S2 and BG3 cells (Kharchenko *et al.* 2011), we queried all chromatin states for an over-representation of associations to PEV. First, we generated an expected proportion of chromatin states using all autosomal variants (Figure S8). We then selected the top 1,000 SMA variants, as rank-ordered by *P*-value, and compared the represented chromatin state proportions to the genome-wide distribution. We found an over-representation of chromatin state c1 using data from both BG3 and S2 cells (> 1 standard deviation) and lowered representation of chromatin state c4, c6 and c7 (< 1 standard deviation). Chromatin state c1 is described as representing active promoter and transcriptional start site (TSS)-proximal regions and consists of regions having H3K4me2, H3K4me3 and H3K9ac histone marks.

Given that chromatin states are correlated with factors that influence function across the genome, we next performed an unbiased query of individual ChIP-chip and ChIP-seq data also made publicly available through the modENCODE project (modENCODE Consortium *et al.* 2010; Kharchenko *et al.* 2011). This dataset is comprised of hundreds of factors sampled across a full range of developmental stages, with a bulk of the data coming from S2 and BG3 cells. Again, for each chromatin factor with prior data, we generated the null distribution of variants expected to exist within each site if randomly drawn from the all autosomes (Figure S9). Variants from the top ranked associations were then compared to expected and assessed for enrichment. We note that several features displayed a strong enrichment with PEV associated variants, including histone modifications, transcription factors and non-transcription factors. We repeated these tests for an over-representation of chromatin state, c1, where sites from all three histone marks, H3K4me2, H3K4me3 and H3K9ac are enriched with top associations (Figure 4). We also observed enrichment for binding sites of know PEV factors, such as JIL-1 (Lerach *et al.* 2006), LSD1 (Di Stefano *et al.* 2007), BEAF-32 (Gilbert *et al.* 2006), among others. Further, we note enrichment of top ranked associations in sites that suggest natural variation has a particular impact on TSS regions that show a “balanced” or “bistable” chromatin state. Bistable chromatin sites are sites that may be influenced to either exhibit active or repressed gene expression. We note statistically significant differences in nearly all bound factors that strongly characterize bistable sites; ASH1, H3K4me1, H3K4me2, H3K4me3, RNA Pol II, including depletion of H3K27me3 (Kharchenko *et al.* 2011). The over-representation of associations in sites bound by known chromatin modifiers suggests that natural autosomal variation interacts with chromatin dynamics and PEV through influencing genome-wide expression rates or chromatin state occupancy and balance, not through altering protein function of individual genic modifiers.

**Figure 4.**
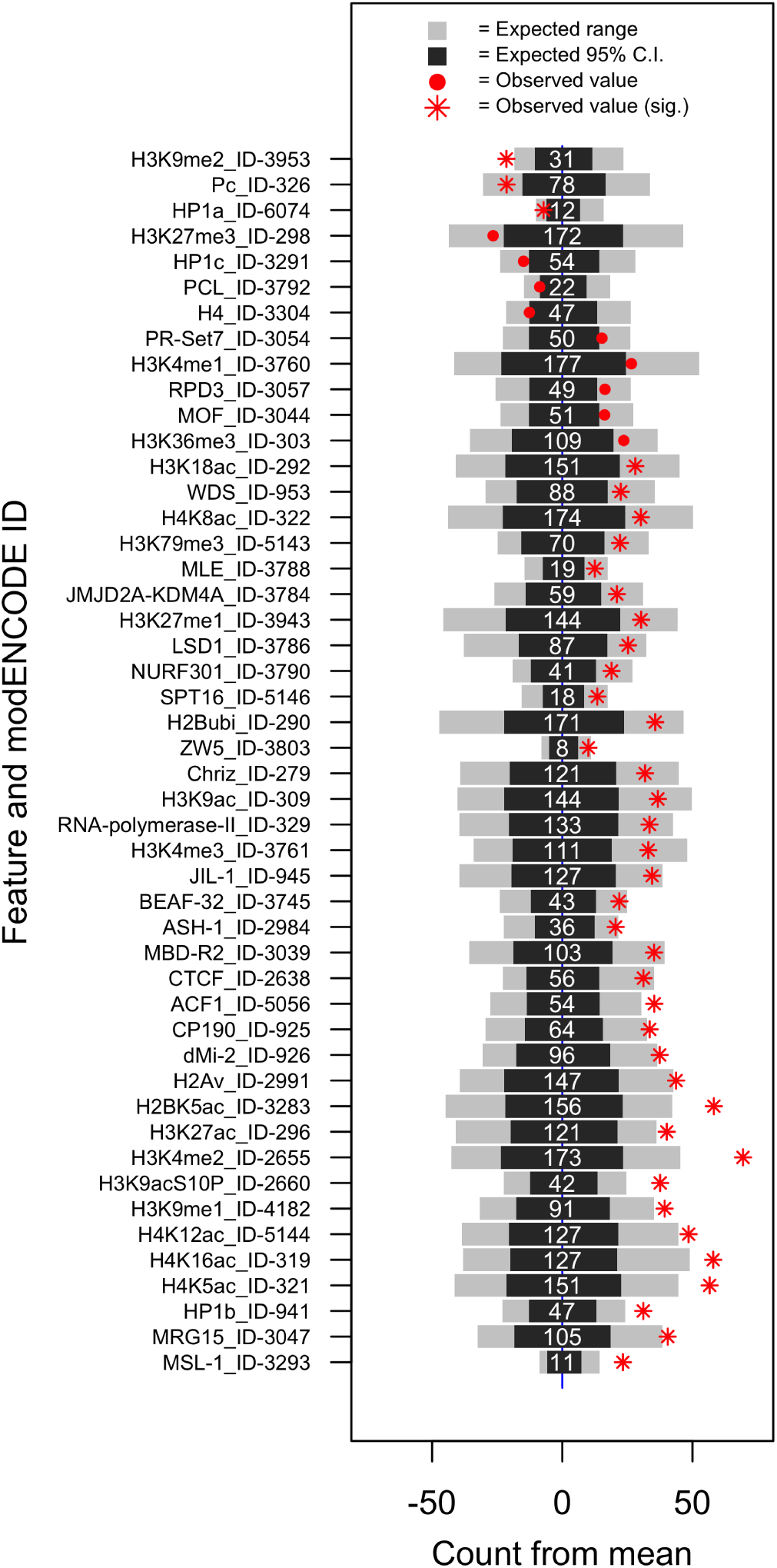
Observed enrichment of variants with association to PEV and chromatin features from S2 cells only. Black bars represent 95% of the expected distribution and gray bars represent the left and right 2.5% tails of the distribution. Numeric values are the average number of variants expected to fall within each chromatin feature. Observed counts from variants within the 1,000 top *P*-value ranked associations are represented in red. Only features with observed values in the 5% tails of the expected distribution are shown. False Discovery Rate (FDR) was used to assess statistical significance across the multiple tests, and significant values are noted.

Remarkably, these findings have precedent in the PEV system and fit extremely well with prior findings indicating a key driver behind mosaic features of variegation is a bistable equilibrium between TF binding and heterochromatin content (Ahmad and Henikoff 2001). Here, it was found that by simply varying levels of a GAL4 transcriptional activator to a heterochromatin-embedded promoter, heterochromatin state could be disrupted. In the model proposed, termed The Site Exposure Model of Variegated Silencing (Widom 1999; Ahmad and Henikoff 2001), kinetics of DNA-histone contact dictate ability of a TF to bind a promoter or enhancer feature, and thus influence gene expression. Features that increase contact between TF binding and activator, include changes to TF abundance or changes to TF binding efficiency such as through mutated underlying binding sequence, changes to nucleosome occupancy (observed through histone and chromatin marks), or changes to abundance of TF guide molecules. Applied to our study, this suggests that each individual shows differences in PEV due to a large number of sites that impact expression rates of factors that then impact binding efficiency of TFs at the *w*^*m4*^ locus. This model also predicts that changes to chromatin content, i.e. an increase or decrease in heterochromatin, can in turn impact sensitive loci throughout the genome and influence gene expression. Indeed, this fits with observations that differing natural Y chromosomes, a giant source of heterochromatin, impact gene expression in autosomes (Lemos *et al.* 2008, 2010).

## Discussion

We designed an assay to identify autosomal non-recessive variants involved in differences in PEV between naturally derived lines of *Drosophila melanogaster*. Despite large PEV-induced pigmentation differences between phenotyped lines, we find little evidence for involvement of polymorphic sites within known Su(var) and E(var) genes that contribute to these differences in PEV. Our top SMA associations further indicate that natural differences in response to PEV are not the primary result of changes in protein function among lines. We instead find that regions of enriched association to PEV are over-represented for promoter and TSS-proximal regions with an additional emphasis on sites that display bi-stable chromatin features. This suggests that autosomal interactions with differences in PEV, i.e. differences in heterochromatin dynamics, are the combined result of many small effect loci that accumulate differences and are linked to modified rates of transcription, either through small changes to specific TF binding sites or broad changes in chromatin state and chromatin mark distributions. Furthermore, our data fit the Site Exposure Model of Variegated Silencing (Ahmad and Henikoff 2001), where a variegated state is the result of bistable features between TF binding and chromatin features. However, as the effect sizes for all associated variants are small, the combined set of variants likely do not fully explain differences in PEV across our sample set; indicating there is yet unobserved genetic variation that accounts for differences among lines. It is important to note that this query of natural variation was not exhaustive and only considered autosomal variants with a particular focus on common variants. We reasoned that a heterozygous screen would be informative, because most Su(var) and E(var) allelic effects are dominant, but we note there is every reason to believe that recessive genetic modifiers of PEV exist also and were missed in our screen. Importantly, we did not query GxG interactions or Y-linked variants, known contributors to PEV and gene expression differences in natural populations (Lemos *et al.* 2008, 2010). Finally, it is important to consider that the autosomal loci identified here only show correlation with differences in PEV; it is not known at this time if these loci represent drivers or are instead the consequence of differences in heterochromatin dynamics.

## Acknowledgments

We thank Dan Barbash and Jason Mezey for thoughtful critique and comments throughout all stages of the work. Several members of the Clark lab provided support through general discussion and expertise, including; Jen Grenier, Clement Chow, Rob Unckless, Julien Ayroles, Angela Early and Grace Chi. This work was supported by R01 GM119125.

---

**Figure S 1.**
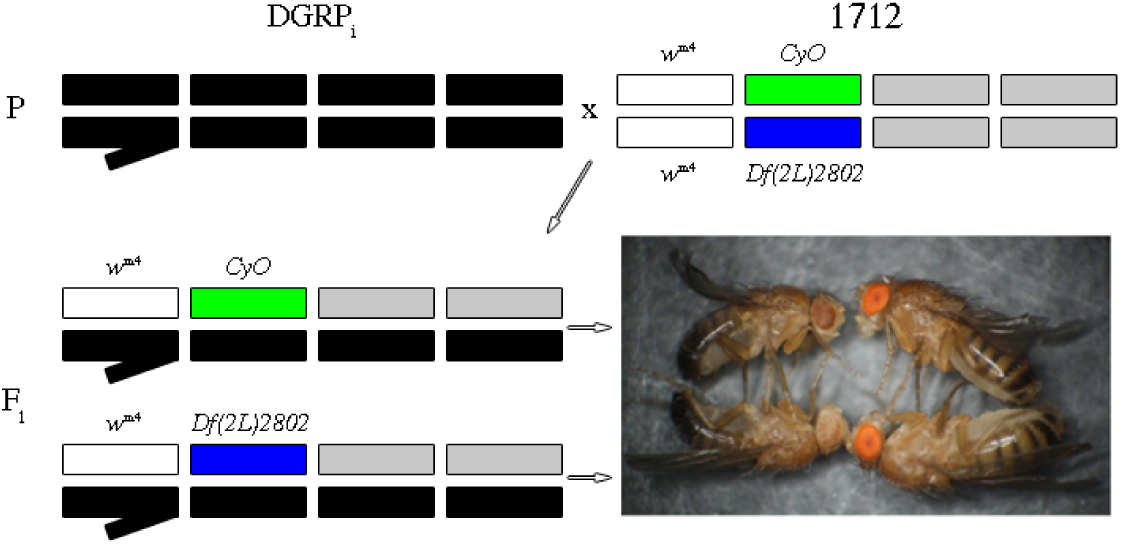
Experimental cross to generate males with unique DGRP haplotypes, *i*, that also carry *w*^*m4*^ and exhibit PEV. F1 males segregate into two variegating classes, those with a *CyO* second chromosome (visible curly wing) and those with *Df(2L)2802* second chromosome (visible straight wing). F1 males were imaged. Females do not display variegation in this cross and were not studied.

**Figure S 2.**
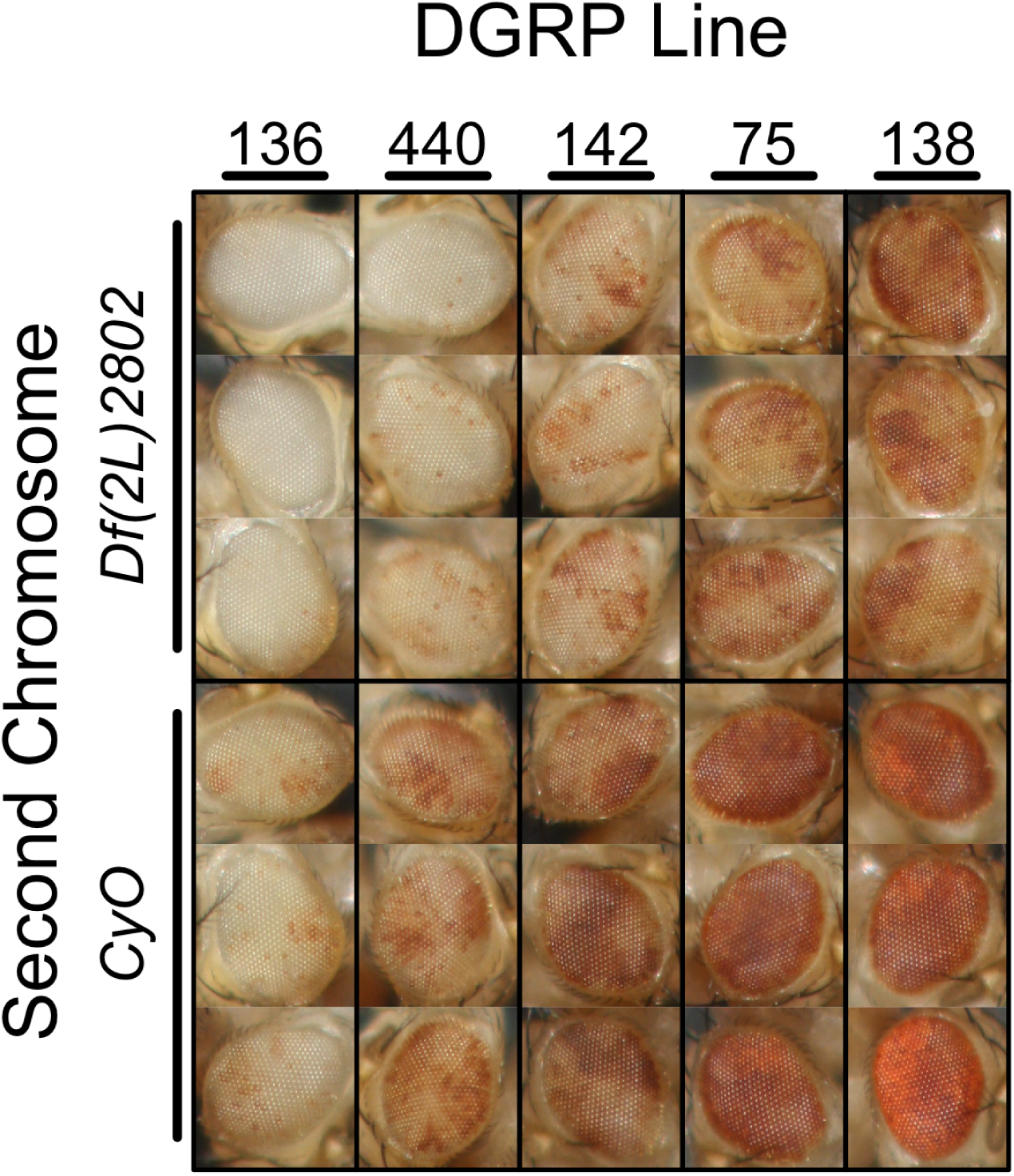
F1 experimental males show a wide range of visible differences in eye pigmentation, spanning a range that lacks pigmentation (DGRP line 136 with a *Df(2L)2802* second chromosome background) to full pigmentation and near-wild-type eyes (DGRP line 138 with a *CyO* second chromosome background. To demonstrate the range of pigmentation in F1 progeny, three replicates are shown from a subset of DGRP line by second chromosome combinations. Even at a gross level, the PEV phenotype shows greater similarities in pigmentation between replicate individuals than between DGRP line or second chromosome background.

**Figure S 3.**
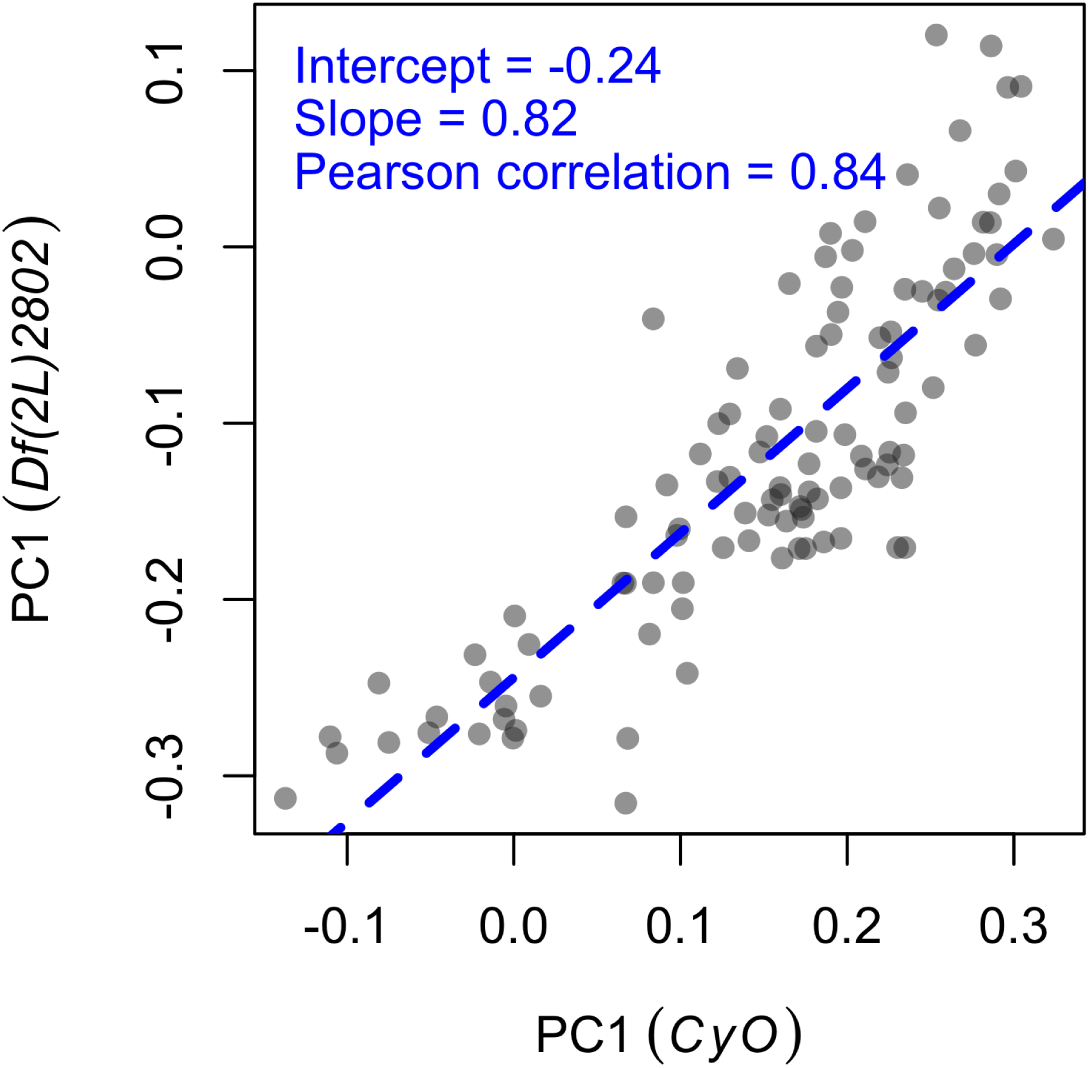
Comparison between PC1 values of PEV populations with *CyO* and *Df(2L)2802* second chromosome backgrounds. Although there is strong correlation in PEV between the two backgrounds, the *CyO* second chromosome shows less variegation (more red eyes and higher PC1 values) with respect to the *Df(2L)2802* second chromosome background.

**Figure S 4.**
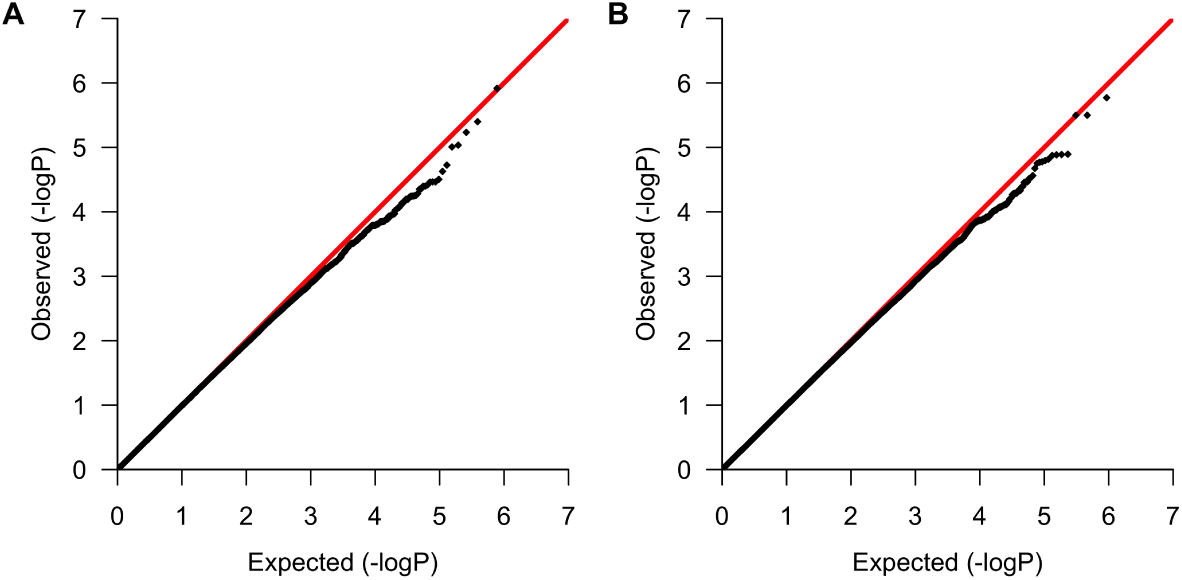
Q-Q plots of expected (red line) and observed (black points) *P*-values from GWAS. **(A)** Experimental population with *CyO* second background. **(B)** Experimental population with *Df(2L)2802* second chromosome background.

**Figure S 5.**
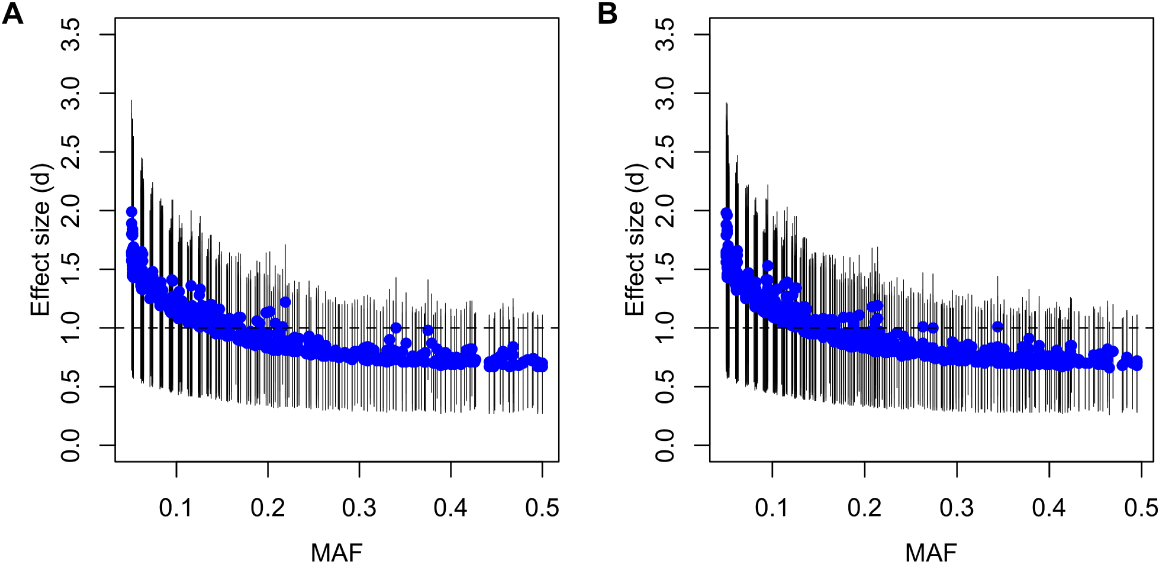
Effect size (Cohen’s *d*) with upper and lower confidence limits of variants having association *P*-values ≤ 0.001. **(A)** Effect sizes of 584 variants from the experimental population with *CyO* second chromosome background. **(B)** Effect sizes of 770 variants from the experimental population with *Df(2L)2802* second chromosome background. 310 and 369 variants from each respective background have effect sizes greater than or equal to 1.

**Figure S 6.**
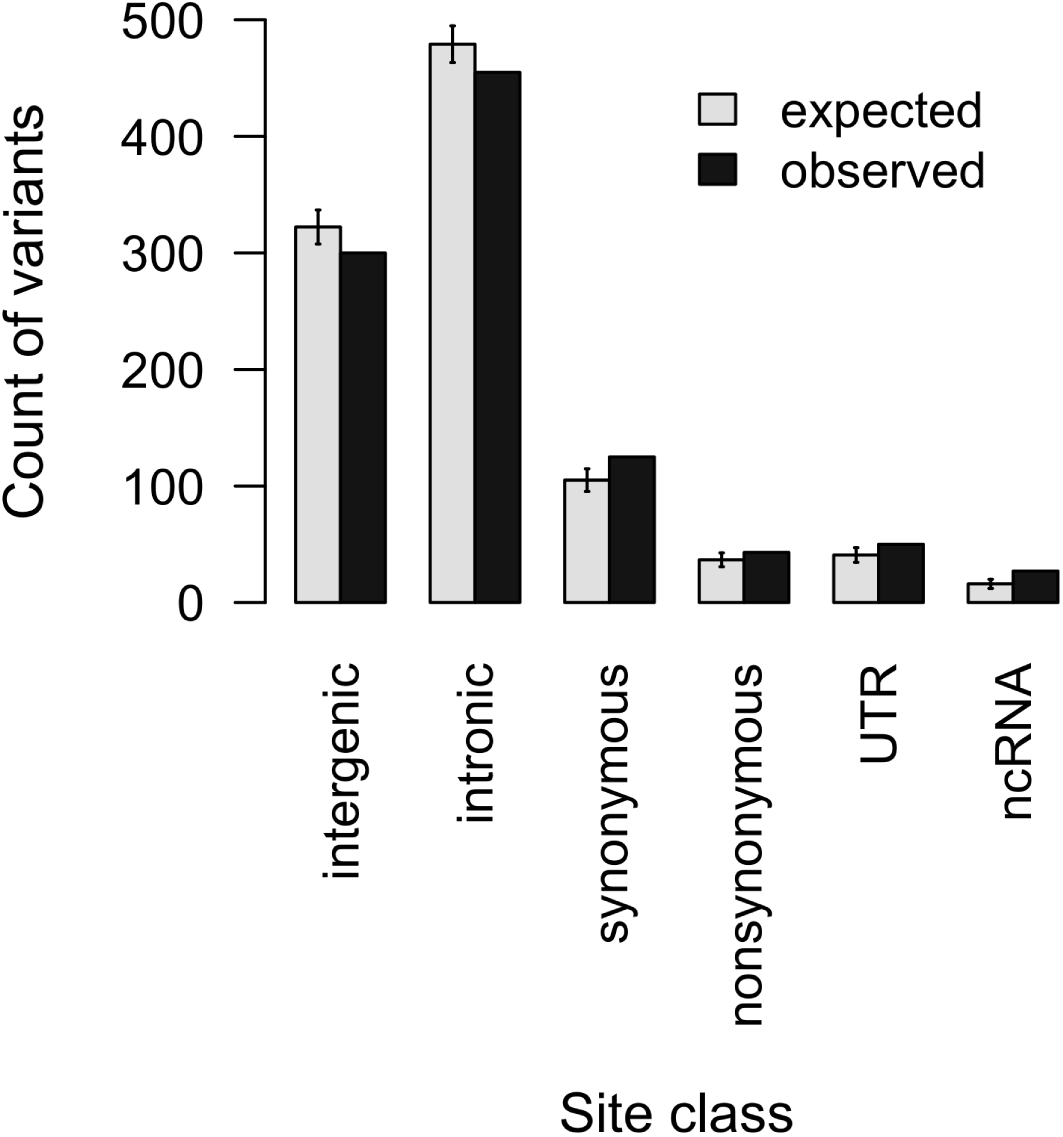
Distribution of site classes between expected counts (average) and observed counts from the 1000 top associated variants. All observed counts show a significant difference from expected (two-sided, one sample *t*-test, Bonferroni corrected *P*-values all ≤ 2.2×10^−16^. Error bars represent the standard deviation across 10,000 randomly drawn samples.

**Figure S 7.**
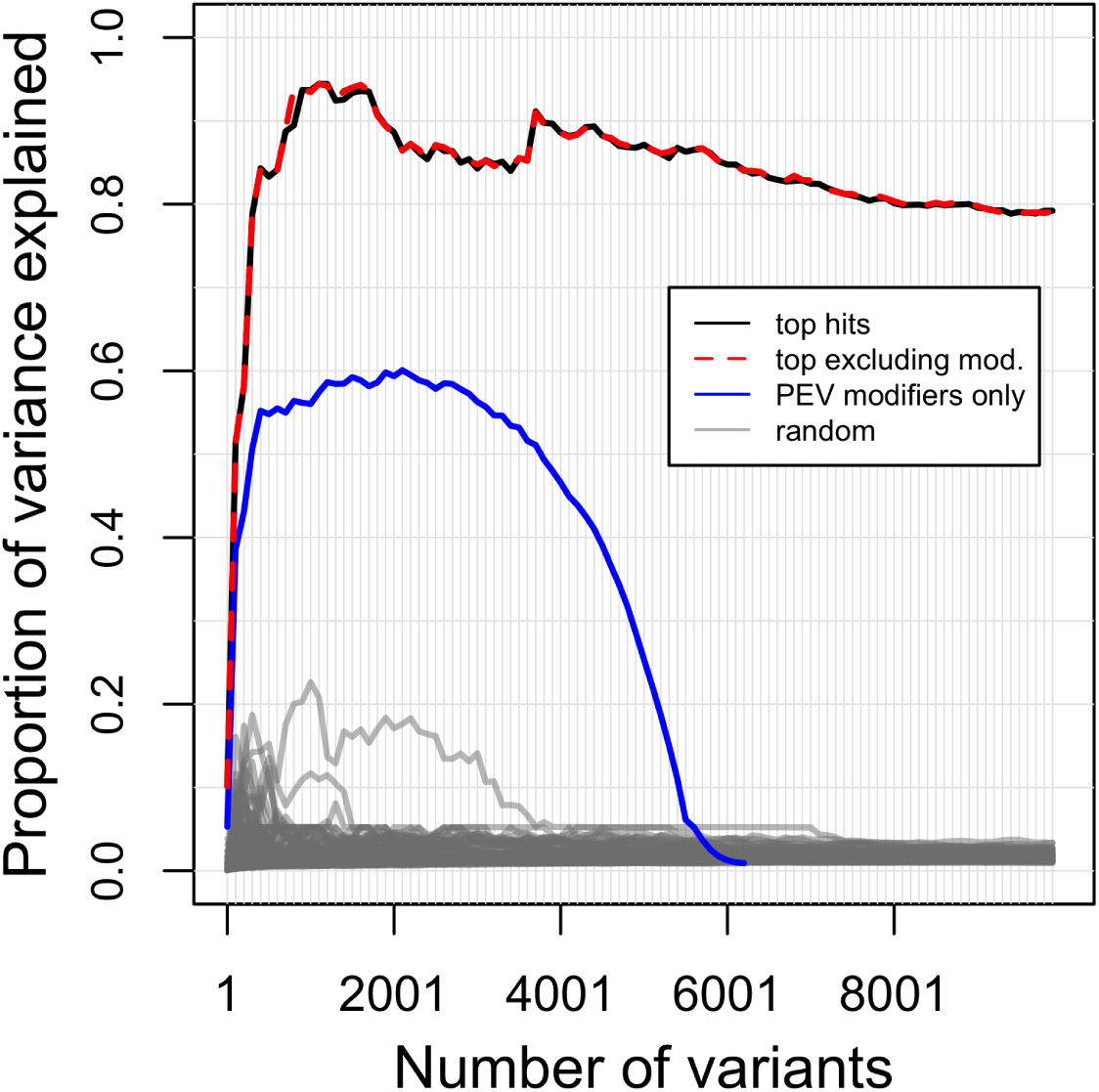
Proportion of between-line phenotypic variance explained within the *Df(2L)2802* second chromosome population using GCTA. Comparison of SNP groupings include; the most significant GWA variants (black line), top hits excluding variants from known PEV genic modifiers (red line), variants within know PEV genic modifiers only (blue line), and randomly selected autosomal variants (gray line).

**Figure S 8.**
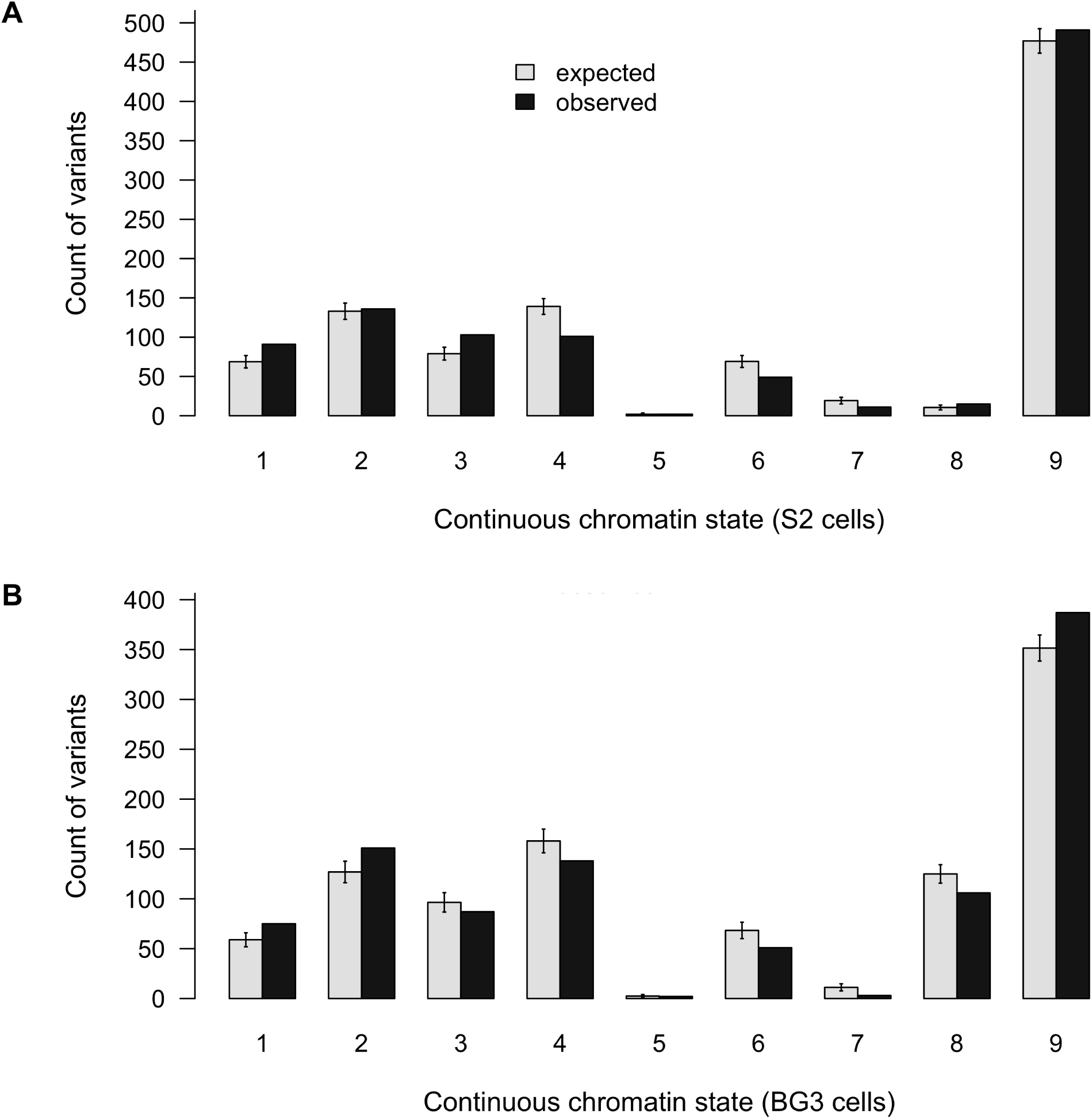
Counts of continuous 9-state chromatin assignments in randomly select autosomal variants (expected) and the top 1,000 *P*-value ranked SMA variants (observed). **(A)** Counts of chromatin state assignments in S2 cells. **(B)** Counts of chromatin state assignments in BG3 cells.

**Figure S 9.**
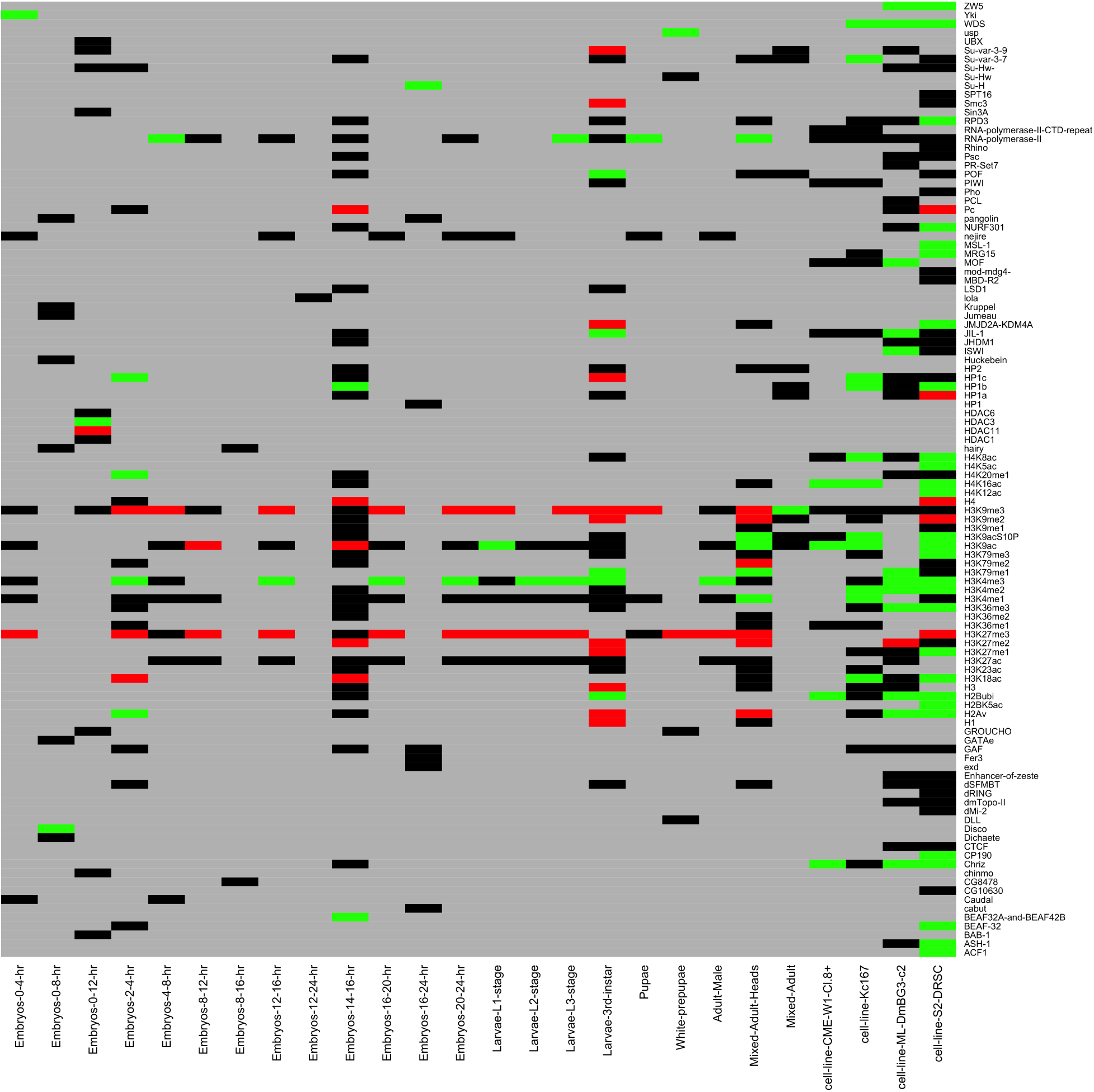
Expected and observed counts of variants within binding sites of chromatin-associating factors (y-axis) and across various developmental stages (x-axis), as defined through ChIP-chip and ChIP-seq binding assays. For each factor and developmental stage, expected distributions were generated through 10,000 iterations of randomly sampling 1,000 variants from all autosomes, and the number of variants residing within marked binding locations were counted. This expected distribution was then compared to the number of variants residing within marked binding locations from the set of 1,000 top ranked associations. Observed counts with respect the expected distribution are as follows; green represents observed counts that are > 97.5% of the expected distribution, red represents observed counts that are < 2.5% of the expected distribution, black represents observed counts that are within 95% of the expected mass, and gray represents missing sample data.

**Table S1.** Heritability of PEV

| Variable | CyO background |  |  |  | Df(2L)2802 background |  |  |  |
| --- | --- | --- | --- | --- | --- | --- | --- | --- |
|  | PC1 | PC2 | PC3 | Adjusted<sup>a</sup> | PC1 | PC2 | PC3 | Adjusted<sup>b</sup> |
| Line | 59.8% | 45.3% | 44.0% | 59.4% | 57.7% | 47.8% | 55.8% | 57.4% |
| Vial | < 0.1% | 0.5% | < 0.1% | < 0.1% | < 0.1% | 0.2% | 0.2% | < 0.1% |
| Residuals | 40.2% | 54.3% | 56.0% | 40.6% | 42.3% | 52.0% | 44.0% | 42.6% |

<sup>a</sup> Adjusted values are calculated by summing across each specific variable as weighted by the proportion of data explained by each PC; 97.4% (PC1), 2.4% (PC2) and 0.2% (PC3). PC3 is not shown above as it explains a negligible amount of data.
<sup>b</sup> See footnote *a*

**Table S2.** GWAS top ranked associations by *P*-value

| Position | Alleles | Gene | Class | CyO background |  |  | *Df(2L)2802* background |  |  |
| --- | --- | --- | --- | --- | --- | --- | --- | --- | --- |
|  |  |  |  | AF | *P*-value | Rank | AF | *P*-value | Rank |
| 2L 12252172 | T/C | *Aret* | Intron | NA | NA | NA | 0.095 | 1.3x10<sup>-5</sup> | 5 |
| 2R 11802982 | C/T | *Sli* | Intron | NA | NA | NA | 0.274 | 1.6x10<sup>-5</sup> | 9 |
| 2R 12408574 | T/C | *Sema-2a* | Intron | 0.126 | 1.8x10<sup>-5</sup> | 6 | 0.124 | 1.7x10<sup>-5</sup> | 10 |
| 2R 18531570 | T/C | NA | Intergenic | 0.051 | 3.1x10<sup>-5</sup> | 8 | 0.05 | 3.5x10<sup>-5</sup> | 19 |
| 3L 1775145 | A/G | *CG13933* | 5'UTR | 0.34 | 5.8x10<sup>-6</sup> | 3 | 0.344 | 3.2x10<sup>-6</sup> | 3 |
| 3L 2081047 | A/G | *sls* | Synonymous | 0.116 | 2.4x10<sup>-5</sup> | 7 | 0.113 | 2.1x10<sup>-5</sup> | 13 |
| 3L 2288054 | G/T | *DmsR-2* | Intron | 0.198 | 9.8x10<sup>-6</sup> | 5 | 0.194 | 1.3x10<sup>-5</sup> | 6 |
| 3L 2568884 | T/C | *msn* | Intron | 0.375 | 4.0x10<sup>-6</sup> | 2 | 0.378 | 1.3x10<sup>-5</sup> | 7 |
| 3L 3595434 | T/G | *Eip63E* | Intron | 0.219 | 1.2x10<sup>-6</sup> | 1 | 0.214 | 1.7x10<sup>-6</sup> | 1 |
| 3L 5909455 | T/C | *CG13288* | Intron | 0.202 | 9.2x10<sup>-6</sup> | 4 | 0.208 | 3.2x10<sup>-6</sup> | 2 |
| 3L 5909456 | G/C | *CG13288* | Intron | 0.208 | 3.5x10<sup>-5</sup> | 11 | 0.214 | 1.3x10<sup>-5</sup> | 4 |
| 3L 12852675 | C/G | *CG32107* | Nonsyn. | 0.333 | 3.4x10<sup>-5</sup> | 9 | 0.337 | 1.2x10<sup>-4</sup> | 78 |
| 3L 12852676 | T/C | *CG32107* | Nonsyn. | 0.333 | 3.4x10<sup>-5</sup> | 10 | 0.337 | 1.2x10<sup>-4</sup> | 79 |
| 3R 4014584 | C/T | NA | Intergenic | NA | NA | NA | 0.116 | 1.5x10<sup>-5</sup> | 8 |

**Table S3.** Known autosomal genic modifiers of PEV

| Chromosome |  |  |  |
| --- | --- | --- | --- |
| 2L | 2R | 3L | 3R |
| *abo* | *Asx* | *AGO2* | *Ace* |
| *aub* | *Atf-2* | *asf1* | *Acf1* |
| *barr* | *BEAF-32* | *ash1* | *Bin1* |
| *CG8677* | *bw* | *brm* | *bon* |
| *Chd1* | *Cap-G* | *E(z)* | *cav* |
| *chm* | *Dcr-2* | *FRG1* | *D1* |
| *dp* | *dom* | *HP4* | *Dcr-1* |
| *Etl1* | *Dp1* | *Jarid2* | *E(var)3-9* |
| *glu* | *E(Pc)* | *JIL-1* | *E2f* |
| *Hel25E* | *egg* | *msl-3* | *Fmr1* |
| *kis* | *LamC* | *not* | *gpp* |
| *lid* | *lat* | *Pc* | *His2Av* |
| *lt* | *Nap1* | *pzg* | *Invadolysin* |
| *lwr* | *Pcl* | *rept* | *jumu* |
| *Mcm10* | *PCNA* | *Rpd3* | *MBD-like* |
| *Mt2* | *Psc* | *RpLP0* | *Mekk1* |
| *piwi* | *psq* | *Sgf11* | *mod* |
| *Psf2* | *SMC2* | *sti* | *mod(mdg4)* |
| *r2d2* | *Su(var)2-10* | *stwl* | *mor* |
| *Sam-S* | *Su(var)2-HP2* | *Su(var)3-3* | *MRG15* |
| *Sir2* | *Su(z)2* | *Su(z)12* | *Orc2* |
| *Sos* |  | *SuUR* | *osa* |
| *Su(var)205* |  | *Trl* | *polybromo* |
| *vig* |  | *Ubp64E* | *Pp1-87B* |
|  |  |  | *pr-set7* |
|  |  |  | *puc* |
|  |  |  | *Rga* |
|  |  |  | *Sce* |
|  |  |  | *Scm* |
|  |  |  | *Snr1* |
|  |  |  | *spn-E* |
|  |  |  | *Su(var)3-7* |
|  |  |  | *Su(var)3-9* |
|  |  |  | *trx* |
|  |  |  | *vig2* |
|  |  |  | *XNP* |

